# Host-like Temperature Unlocks Glucose Uptake and Metabolism in Pathogenic *Leptospira*

**DOI:** 10.64898/2026.07.24.740548

**Authors:** Ryo Ozuru, Michinobu Yoshimura, Deborah A Powers, Takumi Sonoda, Jason A. Papin, Fumiko Obata, Glynis L. Kolling, Kenji Hiromatsu

## Abstract

Pathogenic *Leptospira* transitions between environmental reservoirs and mammalian hosts, expose the bacterium to distinct temperature conditions. Although *Leptospira* is generally considered to rely its growth primarily on long-chain fatty acids and to have limited capacity to use glucose, whether host-like temperature alters glucose-associated metabolic capacity remains unclear. Here, we used transcriptome-integrated genome-scale metabolic modeling to examine temperature-dependent metabolic states in pathogenic *Leptospira*. Models contextualized with transcriptomic data from cultures at 37°C predicted increased flux through glucose transport, glucose phosphorylation, and downstream glucose-associated reactions than it did at 29–30°C. Similar temperature-dependent predictions were obtained for another pathogenic *Leptospira* species under *in vitro* conditions. Consistent with these findings, analysis of leptospires cultivated intraperitoneally in dialysis membrane chambers also predicted enhanced glucose uptake and metabolism under host-temperature conditions. *In vitro* validation experiments showed increased bacterial uptake or accumulation of a fluorescent glucose analogue, elevated expression of candidate glucose transporter genes, and glucose-dependent enhancement of bacterial growth at 37°C. Treatment with a glucose degradation pathway inhibitor, 2-deoxy-D-glucose further supported a contribution of glucose-associated processes to proliferation at 37°C. Together, these findings indicate that pathogenic *Leptospira* displays condition-dependent glucose uptake and glucose-associated metabolic activity that become apparent at host-like temperature, opposed to glucose-incompetent in an environmental-like temperature. This prediction-driven framework refines the conventional view of *Leptospira* carbon metabolism and provides a basis for future studies of glucose-associated metabolic capacity during mammalian infection.

**Importance:** Environmental reservoirs play a central role in the transmission of many bacterial pathogens, yet the metabolic mechanisms that enable adaptation to both environmental and host-associated niches remain largely unknown. Using a prediction-driven framework that combines contextualized genome-scale metabolic modeling with multi-omics analyses and experimental validation, we discovered that pathogenic *Leptospira* activates glucose uptake and metabolism only under host-like conditions. This finding resolves a long-standing misconception arising from studies performed under conventional laboratory conditions and fundamentally revises our understanding of *Leptospira* physiology. More importantly, our study establishes an integrative systems biology strategy for uncovering condition-dependent metabolic traits that would be difficult to identify experimentally alone, with broad applicability to diverse environmentally transmitted pathogens.

## Introduction

The zoonotic spirochete *Leptospira* can persist outside hosts and survive under various environmental conditions. While *Leptospira* proliferates most efficiently when cultured at 30°C (1), pathogenic species also encounter and tolerate the higher body temperature of mammalian hosts, typically around 37°C. Growth differences between these two temperatures indicate that the physiological and metabolic state of *Leptospira* under host-like temperature conditions may differ from that of under optimal environment-mimicking culture conditions. Such differences are likely shaped, at least in part, by the availability of carbon sources and by the metabolic capabilities of *Leptospira*. Indeed, numerous studies have demonstrated temperature-dependent changes in *Leptospira* transcript and protein expression (2–8), whereas relatively few have focused specifically on metabolism. As a result, higher temperature-induced reductions in growth rate are often interpreted as a general decrease in metabolic activity without detailed analysis of the underlying metabolic pathways. Elucidating these temperature-associated metabolic changes may help identify condition-dependent biomarkers and prioritize metabolic processes relevant to severe leptospirosis.

Defining the carbon sources of microorganisms is fundamental to understanding their ecology, host specificity, and physiological responses to environmental and host-related conditions. A key metabolic feature of *Leptospira* is its dependence on long-chain fatty acids as major carbon and energy sources (9, 10). This metabolic feature has been proposed to contribute to *Leptospira* persistence and proliferation in host tissues, including lipid-rich niches (11), although the context-dependent use of alternative carbon sources remains less well understood. Interestingly, while glucose is commonly utilized as a carbon source by many microbes, the zoonotic spirochete *Leptospira* has long been regarded as a notable exception. This bacterium lacks several enzymes involved in the glycolytic pathway, which is the initial route for glucose catabolism (12). Moreover, *Leptospira* exhibits extremely low expression of glucose transporters (13), resulting in minimal intracellular glucose levels (14). Despite possessing several glycolytic enzymes and a potential for gluconeogenesis, *Leptospira*’s carbohydrate metabolism remains poorly characterized.

Genome-scale metabolic network reconstructions (GENREs) have emerged as powerful tools for investigating microbial metabolism and predicting condition-dependent metabolic capabilities (15). By enabling the prediction of metabolic fluxes and the simulation of systems-level metabolic behavior, GENREs facilitate metabolic engineering and provide mechanistic hypotheses for microbial physiology (15, 16). In the context of infectious diseases, GENREs have been employed to prioritize antimicrobial targets (17) and to explore pathogen metabolism (18, 19). These models have also been used to study pathogen behavior including the evolution of antibiotic resistance, the production of virulence factors, and host-pathogen interactions (20). For example, GENRE-based analyses of *Pseudomonas aeruginosa* have revealed key metabolic adaptations during chronic lung infections in cystic fibrosis patients, shedding light on disease progression and therapeutic vulnerabilities (21). Although GENREs have previously been constructed for *Leptospira* (22), these models have yet to be fully leveraged to examine its temperature-dependent metabolic responses under host-like conditions.

In this study, we constructed GENREs of pathogenic *Leptospira* to investigate how its metabolic network responds to changes in external temperature. We integrated publicly available transcriptomic datasets derived from cultures grown at different temperatures and from host-like conditions into the models. This approach enabled us to contextualize the *Leptospira* metabolic model and to generate condition-specific predictions that better reflect transcriptomic states under physiologically relevant conditions. Using this framework, we aimed to identify model-predicted metabolic changes associated with temperature shifts and host-associated environments. Furthermore, we experimentally examined the relationship between these predicted metabolic changes, particularly those related to glucose-associated metabolism, and physiological phenotypes in *Leptospira* including glucose uptake and growth dynamics (Fig. 1). Through this integrative approach, we sought to evaluate whether GENREs can uncover condition-dependent metabolic capacities in pathogenic *Leptospira*.

**FIG 1.**
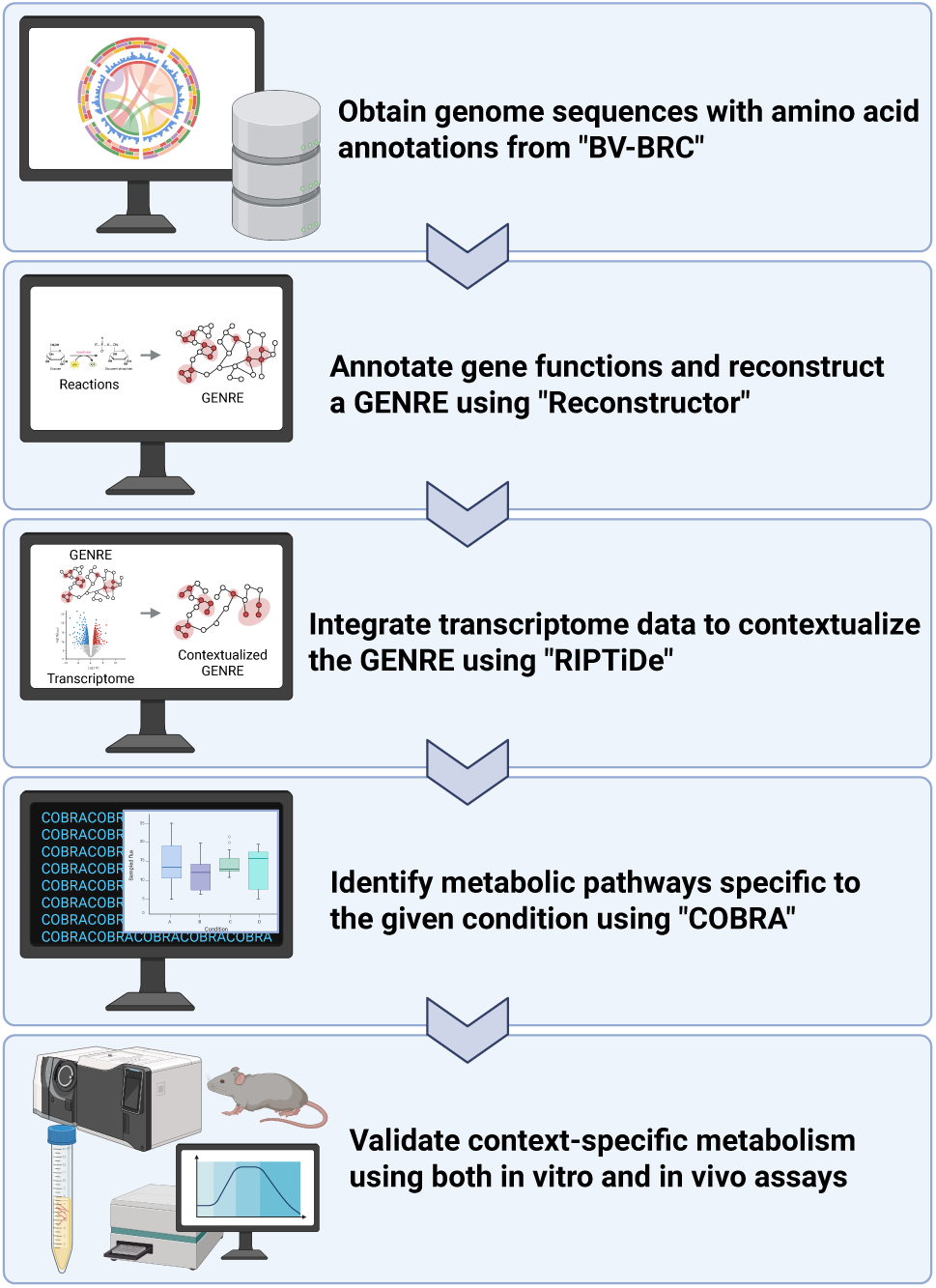
Workflow for genome-scale metabolic network reconstruction, context-specific analysis and validation. Created in BioRender. Ozuru, R. (2026) https://BioRender.com/k7boo1a

## Results

### Temperature-dependent metabolic changes predicted by transcriptome-integrated GENREs of pathogenic *Leptospira*

Draft GENREs of pathogenic *Leptospira* were constructed using Reconstructor (23), a tool that automatically builds draft metabolic models based on the amino acid– annotated FASTA files. The accession numbers of the genome data used, along with genome size, gene count, and model characteristics—such as the number of genes, reactions, and metabolites, as well as MEMOTE (24) scores——are summarized in Table S1. The MEMOTE score is a comprehensive quality metric that assesses the structural completeness, annotation quality, consistency, and simulation readiness of genome-scale metabolic models. To refine these models and generate condition-specific metabolic predictions, we integrated publicly available transcriptomic data from the NCBI Gene Expression Omnibus (GEO). Specifically, we used dataset GSE168507 (25), which contains transcriptomic profiles of the highly pathogenic *L. borgpetersenii* strain JB197 cultured at 29 °C and 37 °C. Data integration was performed using RIPTiDe (26), an algorithm that incorporates transcriptomic data to inform reaction activity and flux distributions within GENREs. Flux balance analysis (FBA) of the context-specific models predicted reduced growth potential at 37 °C compared to 29 °C (Fig. 2A), that the data is consistent with the known optimal growth temperature of *Leptospira* around 30 °C. Dimensionality reduction of the predicted reaction flux profiles separated the two temperature conditions (Fig. 2B). To identify reactions that contributed most to this temperature-associated separation, we applied a random forest classifier, which highlighted key metabolic reactions distinguishing the two context-specific models (Fig. 2C, D and Fig. S1).

**FIG 2.**
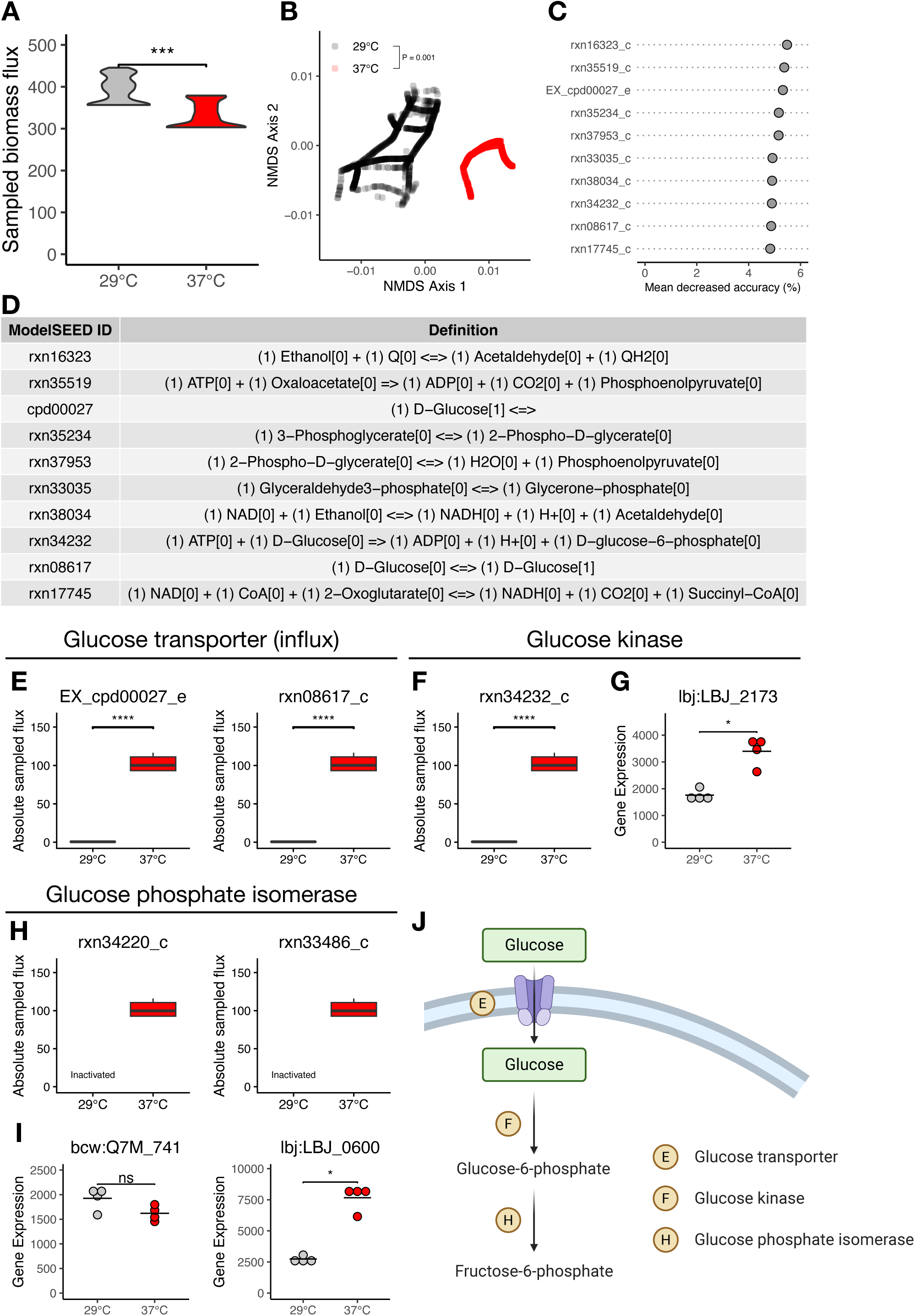
Comparison of metabolic characteristics of contextualized GENREs under different temperature conditions. **A**, Biomass levels at each temperature condition. **B**, Comparison of reaction flux distributions for metabolic reactions shared across all temperature conditions. **C**, List of metabolic reactions in reaction IDs that distinguish temperature conditions, was identified using a machine learning approach. Reaction IDs are from the ModelSEED database. All the actual flux plots (29°C vs. 37°C) are shown in Extended Data Fig. 1. **D**, Reaction IDs and definitions listed in **c**. (n) indicates the number of molecules involved in the reaction. Compartments [0] and [1] represent the intracellular and extracellular spaces, respectively. **E,F**, Flux balance analysis results for selected reactions at different temperatures from **c** and Extended Data Fig. 1. Each plot title corresponds to a reaction ID from the ModelSEED database. **G**, Expression of the gene associated with rxn34232_c. **H**, Flux balance analysis results for selected reactions that were inactive at 29 °C but became active at higher temperatures. **I**, Expressions of the genes associated with rxn34220_c and rxn33486_c. **J**, Overview of glucose metabolism in *Leptospira*. Panels **E**, **F** and **H** correspond to specific reactions indicated in this pathway. Created in BioRender. Ozuru, R. (2026) https://BioRender.com/lwavfld Statistical significance was evaluated using the Wilcoxon rank-sum test (**A**, **G** and **I**), the Dunn test with Bonferroni correction (**E** and **F**) and permutational ANOVA (permANOVA) (**B**). *P < 0.05, \*\**p* < 0.01, \*\*\**p* < 0.001, and \*\*\*\**p* < 0.0001

### Model-predicted activation of glucose-associated reactions under temperature upshift in *Leptospira*

Because *Leptospira* exhibits reduced growth at 37 °C (Fig. 2A), it is expected that the magnitude of metabolic fluxes also decreases under this condition. Nevertheless, only three reactions (glucose transporter; EX_cpd00027_e and rxn08617_c, and glucose kinase; rxn34232_c) showed an increase in absolute flux values at 37 °C, and we therefore focused on these reactions (Fig. 2E and F). The glucose kinase reaction (rxn34232_c) corresponds to the gene ID lbj:LBJ_2173 in the model, which also exhibited increased expression at 37 °C according to the original RNA-seq data (Fig. 2G). In addition, we examined reactions that were removed at 29°C but retained at 37 °C during RIPTiDe contextualization of both models (Fig. 2H). These reactions were associated with glucose phosphate isomerase, which converts glucose 6-phosphate to fructose 6-phosphate (Fig. 2J reaction H) with two steps, mediated by rxn34220 and rxn33486, corresponding to the conversion to beta-glucose 6-phosphate and then to fructose 6-phosphate, respectively (Fig. 2H). Both reactions are associated with bcw:Q7M_741 or lbj:LBJ_0600 genes in the model, but only lbj:LBJ_0600 showed increased expression at 37°C in the original RNA-seq data (Fig. 2I). Together, these findings indicate that transcriptome-integrated model predicts a 37 °C-associated increase in glucose uptake, phosphorylation, and downstream conversion capacity toward fructose-6-phosphate in pathogenic *Leptospira* (Fig. 2J).

### Predicted glucose-associated metabolic changes in additional *Leptospira* species *in vitro* and under host-mimicking conditions

To determine whether the model-predicted glucose-associated changes were specific to *L. borgpetersenii* strain JB197, we conducted a similar analysis for two additional *Leptospira* species using publicly available transcriptomic datasets along with the species’ corresponding metabolic models. First, we integrated transcriptomic data from *L. interrogans* serovar Manilae strain L495 cultured at 30 °C and 37 °C (GSE92976 (27)) into its metabolic model and compared the predicted flux distributions (Fig. S2). This analysis suggested a temperature-associated pattern similar to that predicted for JB197. Next, we analyzed transcriptomic data from a dialysis membrane chamber (DMC) model (GSE53818 (28)), in which *L. interrogans* serovar Copenhageni was encapsulated in a dialysis membrane and implanted into the rat peritoneal cavity to mimic *in vivo* infection. A similar glucose-associated pattern was also predicted under this host-like conditions (Fig. S3, DMC vs. *in vitro* (IV)). These findings suggest that predicted glucose-associated metabolic changes are not limited to *L. borgpetersenii* JB197, but may be shared across the pathogenic *Leptospira* species within host-like conditions examined here.

### Temperature-dependent glucose uptake and glucose-supported growth by ***Leptospira* at 37 °C**

*Leptospira* is generally believed to rely primarily on long-chain fatty acids as carbon sources, with limited capacity to uptake and metabolize glucose. To re-evaluate this assumption, we investigated glucose uptake at different incubation temperatures using a fluorescence-labeled glucose analogue (Fig. 3A). Notably, *Leptospira* exhibited significantly greater uptake of fluorescent glucose at 37 °C compared to 30 °C. This uptake was competitively inhibited by the addition of 10% unlabeled glucose, supporting the specificity of the transporter-mediated process. Based on the hypothesis that increased glucose uptake would be accompanied by upregulation of the corresponding transporter genes, we quantified the expression of previously reported glucose transporter genes (13) by RT-qPCR (Fig. 3B). At 48 h, the expression of LA2133 and LB245 was significantly elevated at 37°C compared with that at 30°C. To assess whether glucose availability affects bacterial proliferation in temperature-dependent manner, we supplemented EMJH medium with glucose and compared growth curves at 30°C and 37 °C (Figs. 3C, D). At 30 °C, glucose supplementation had no significant effect on proliferation. In contrast, at 37 °C, proliferation increased with glucose concentrations up to 2.5 mM. These results physiologically support the model prediction that host-like temperature is associated with an increase in glucose uptake capacity and glucose-supported growth in *Leptospira*.

**FIG 3.**
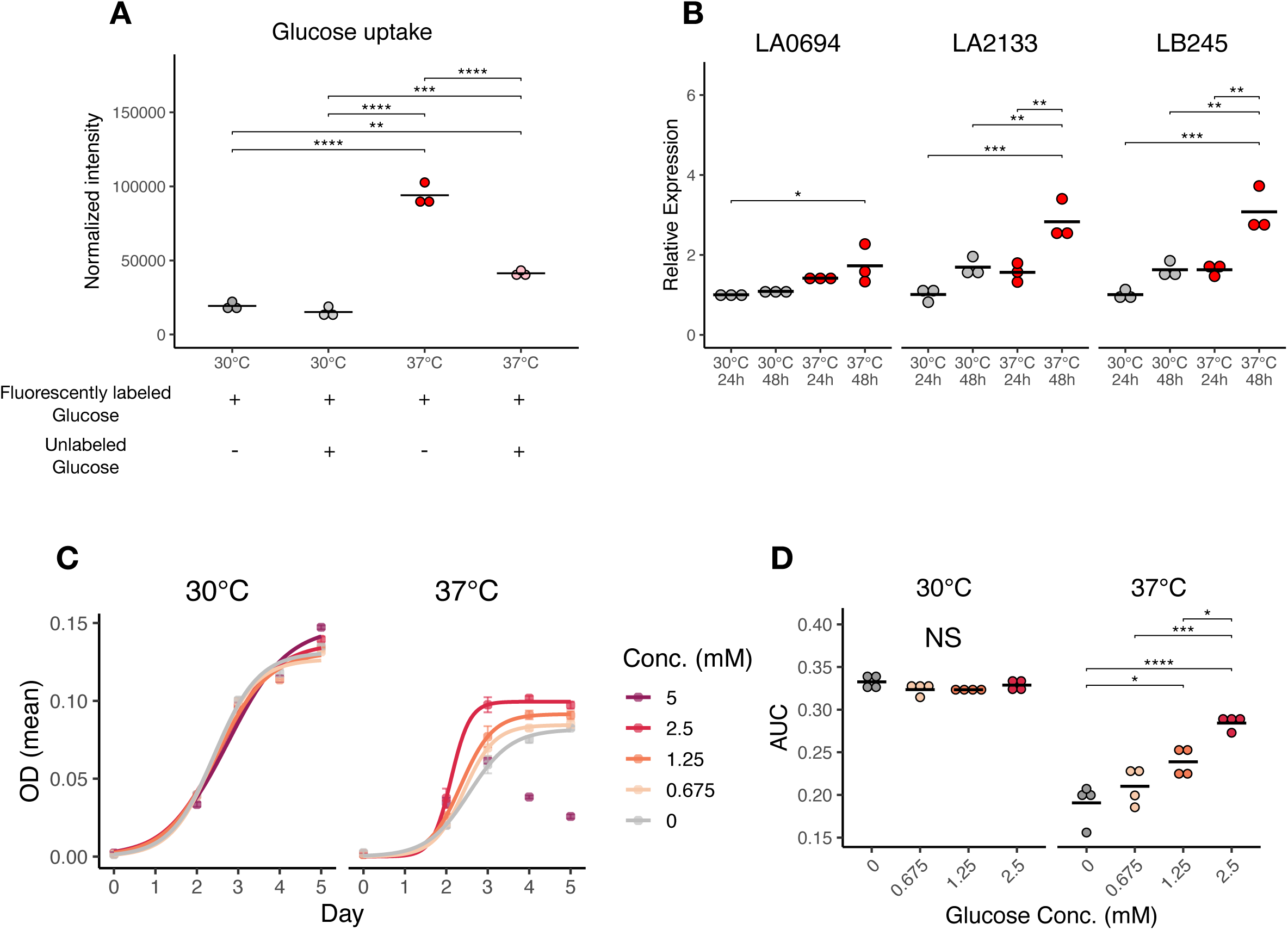
Comparison of glucose uptake and metabolic capacity of *Leptospira* at different temperatures. **A**, Glucose uptake capacity at various temperatures, normalized by bacterial cell number. Competition assays were performed by co-incubating fluorescently labeled glucose with unlabeled glucose. **B**, Expression levels of known glucose transporter genes under 30°C vs. 37°C. **C**, Growth curves of *Leptospira* in EMJH medium supplemented with glucose. **D**, The area under the curve (AUC) from **C**. Statistical significance was evaluated using the one-way ANOVA followed by Tukey’s HSD test. *P < 0.05, \*\**p* < 0.01, \*\*\**p* < 0.001, and \*\*\*\**p* < 0.0001

### Contribution of glucose-associated metabolic activity to *Leptospira* proliferation under host-like temperature

Fructose 6-phosphate, a model-predicted downstream product of glucose-associated reactions, serves as a precursor for the pentose phosphate pathway and the aromatic amino acid biosynthesis pathway (Fig. 4A). These pathways are key metabolic branches that provide reducing power, nucleotides, and precursors for protein and coenzyme synthesis, all of which support bacterial growth. We therefore hypothesized that perturbation of this metabolic node would affect *Leptospira* proliferation, particularly under conditions in which glucose-associated reactions are predicted to be more active. To test this hypothesis, we performed *in vitro* growth assays in the presence of a pathway inhibitor: 2-deoxy-D-glucose (2DG), a glucose analogue that can interfere with glycolysis (Fig. 4A). Although growth inhibition by 2DG at 60 µM was also observed at 30 °C (Figs. 4B and C), the reduction in proliferation was more pronounced at 37 °C (Fig. 4D). These findings support the idea that glucose-associated metabolic activity contributes to pathogenic *Leptospira* proliferation under host-like temperature conditions. Together, these data are consistent with a temperature-dependent contribution of glucose-associated metabolic activity to pathogenic *Leptospira* proliferation *in vitro*.

**FIG 4.**
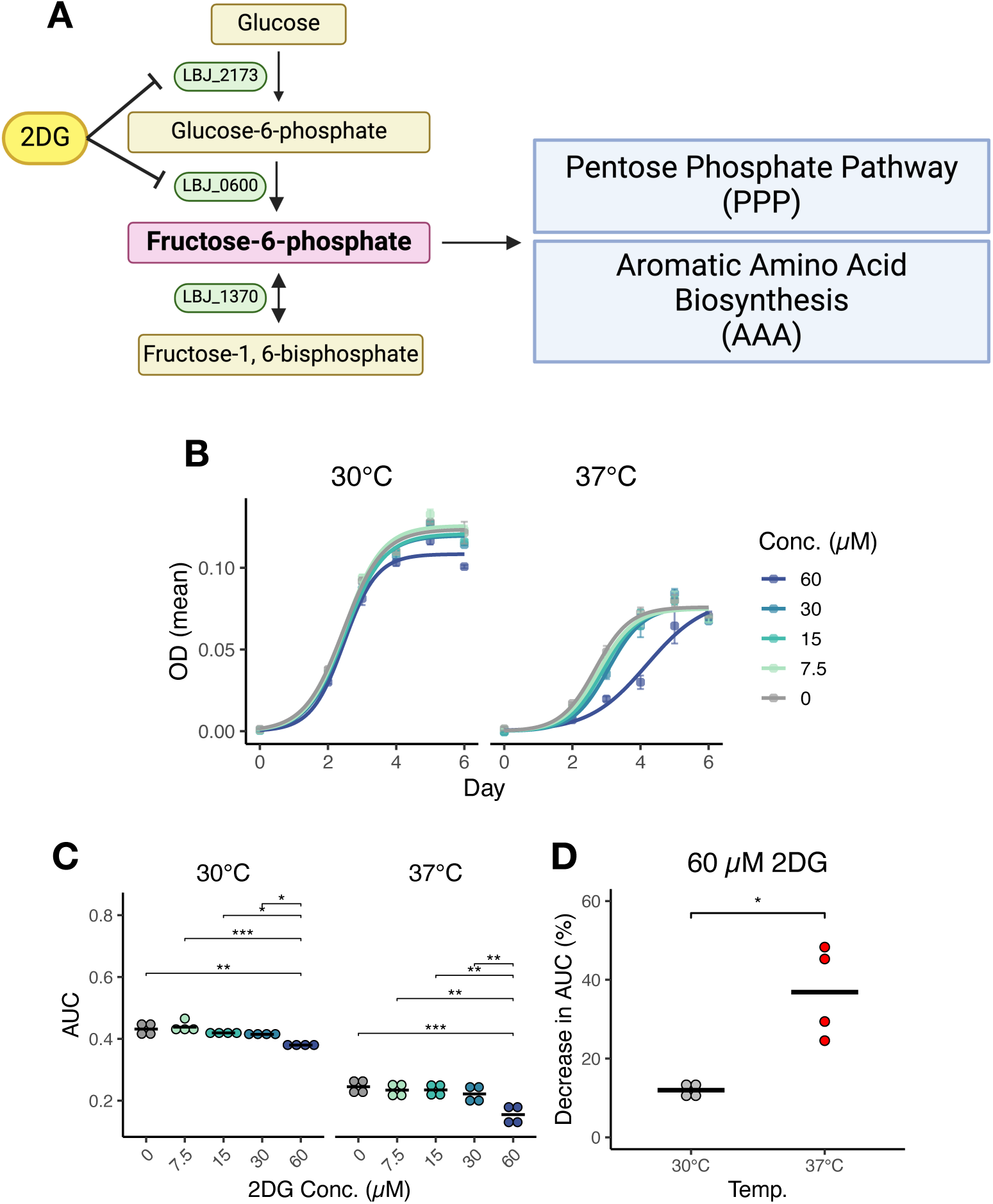
Inhibition of *Leptospira* growth at different temperatures by targeting glycolytic and gluconeogenic pathways. **A**, Schematic diagram of glycolytic pathways active at 37°C. Fructose 6-phosphate, which is intensively synthesized under these conditions, is expected to serve as a precursor for the pentose phosphate pathway (PPP) and aromatic amino acid (AAA) biosynthesis. Enzymes of *L. borgpetersenii* strain JB197 catalyzing each reaction are shown in green (LBJ_XXXX). Created in BioRender. Ozuru, R. (2026) https://BioRender.com/rg4lq23 **B**, Growth curves of *Leptospira* at different temperatures in EMJH medium supplemented with 2-deoxyglucose (2DG). **C**, The area under the curve (AUC) from **B**. **D**, Decrease in the AUC. The AUC values without 2DG at each temperature were normalized to 100%. Statistical significance was evaluated using the one-way ANOVA followed by Tukey’s HSD test (**C**) and Wilcoxon’s Rank Sum test (**D**). *P < 0.05, \*\**p* < 0.01, \*\*\**p* < 0.001, and \*\*\*\**p* < 0.0001

## Discussion

*Leptospira* has distinctive nutrient requirements that differ markedly from those of most other bacteria. It has long been considered to rely primarily on long-chain fatty acids and to have limited capacity to use glucose as a carbon source. In addition, *Leptospira* grows optimally at 30 °C, a temperature lower than that of its mammalian hosts. Accordingly, many previous studies have employed culture conditions based on these established growth characteristics. However, as a pathogen that transitions between environmental reservoirs and host organisms, *Leptospira* is likely to adjust its physiological and metabolic state in response to changes in external conditions, including temperature.

In this study, we provide evidence that pathogenic *Leptospira* displays temperature-dependent glucose uptake, glucose-associated metabolic activity, and glucose-supported growth when cultured at elevated temperatures (Fig. 5). This temperature-dependent glucose-associated phenotype may contribute to growth under host-like conditions (Fig. S3). Most previous investigations assessing *Leptospira* metabolism, including those focused on carbon source utilization, were conducted at 30 °C, the optimal *in vitro* growth temperature. Our findings highlight the importance of considering physiological parameters, such as host body temperature, when interpreting *Leptospira*’s metabolic behavior. Careful attention to experimental conditions is essential when extrapolating *in vitro* metabolic observations to host-associated contexts.

**FIG 5.**
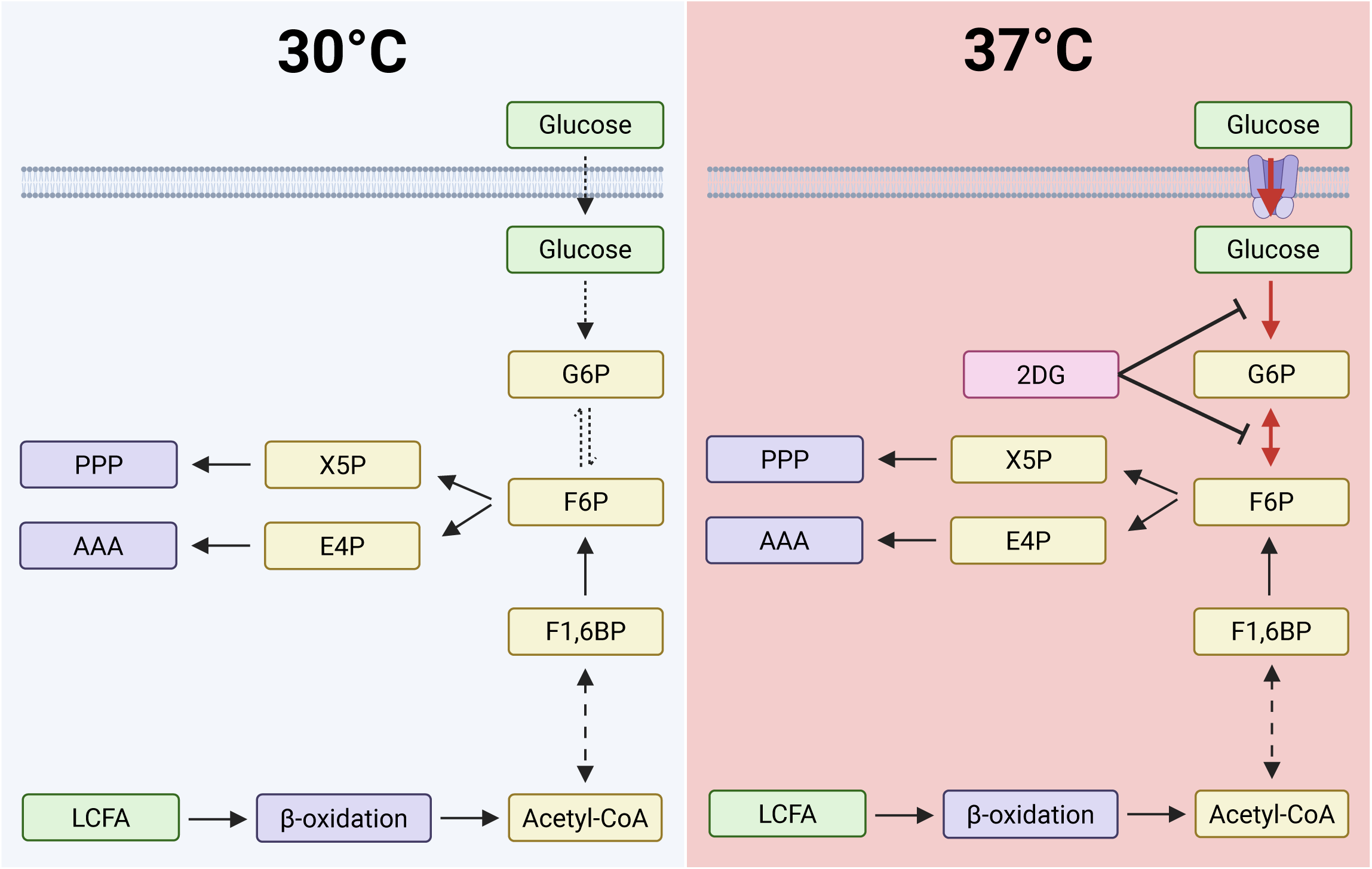
Graphical summary of this study. **Left**, At 30°C, the optimal temperature for *Leptospira* proliferation, glucose uptake and metabolism are inactive. Instead, pyruvate is synthesized from alternative carbon sources such as glycerol for energy production. **Right**, At 37°C, glucose uptake and its conversion to fructose 6-phosphate (F6P) are enhanced. The resulting F6P is channeled into the pentose phosphate pathway (PPP) and aromatic amino acid (AAA) biosynthesis to produce bacterial components. Inhibition of glycolytic pathways by 2DG impairs *Leptospira* growth. Created in BioRender. Ozuru, R. (2026) https://BioRender.com/qp9tha8

This study provides new insights into the temperature responsiveness of glucose-associated metabolism in pathogenic *Leptospira*. Previous genome analysis of *L. interrogans* serovar Copenhageni indicated that this organism possesses nearly all the enzymes comprising the glycolytic pathway (29). The predicted pathway includes a pyrophosphate-dependent fructose-6-phosphate 1-phosphotransferase instead of the conventional ATP-dependent phosphofructokinase, while the initial phosphorylation of glucose is catalyzed by glucokinase. The authors of the genome analysis therefore concluded that the limited utilization of glucose was more likely attributable to constraints in glucose transport than to deficiencies in the downstream metabolic pathway. This interpretation is consistent with our finding that glucose transporter encoding genes LA2133 and LB245, referred to as *putP*-like genes (13) and distinct from LA0694 (which encodes the transporter LIC12908 described in the previous study (29)), exhibited increased expression at host-like temperature (Fig. 3B). Increased expression of these genes may contribute to enhanced glucose uptake (Fig. 3A), alongside the temperature-associated upregulation of genes encoding glucose-associated metabolic enzymes (Fig. 2G and H). Although *putP* is generally annotated as a proline-sodium symporter, homologs in another bacterium have been shown to transport glucose (30). It is therefore possible that the *putP*-like genes in *Leptospira* contribute to condition-dependent glucose uptake, although their substrate specificity and transport activity remain to be experimentally validated.

The question of whether *Leptospira* utilizes glucose as a carbon source has been debated since the organism’s initial discovery. In the late 1910s, Noguchi reported that the addition of glucose to the culture medium did not affect the growth of *Leptospira* (31), leading to the long-standing view that the bacterium has limited capacity to utilize glucose. In the 1960s, Ellinghausen demonstrated that glucose supplementation significantly stimulated the growth of a *Leptospira* isolate obtained from frog kidney tissue, particularly when combined with polysorbate 80 in the culture medium (32). However, the isolate used in that study originated from an amphibian host (*Rana pipiens*), a poikilothermic organism with a physiological temperature range markedly different from that of mammalian hosts. Accordingly, the metabolic characteristics of this isolate, including its temperature preferences and nutrient dependencies, may not fully represent those of pathogenic *Leptospira* strains isolated from homeothermic animals. Indeed, Ellinghausen observed that growth was optimal at 29 °C and severely impaired at 34 °C. In our study, we found that candidate glucose transporter gene expression, glucose uptake and glucose-associated metabolic activity increased in a mammalian-pathogenic *Leptospira* at 37 °C, a temperature that better approximates mammalian host body temperature. These findings suggest that temperature may modulate glucose uptake and glucose-associated metabolic activity, potentially through the induction of specific transporter systems or metabolic enzymes. While Ellinghausen’s results support the possibility that some *Leptospira* isolates can benefit from glucose supplementation, they did not address whether this capacity is modulated by temperature in mammalian-pathogenic strains. These historical and experimental observations shift the question from whether *Leptospira* can use glucose to when and under which physiological conditions glucose-associated metabolic capacity becomes apparent.

Several studies have examined aspects of glucose-associated physiology in *Leptospira*. Yanagihara *et al.* characterized glucose-6-phosphate isomerase, which is one of the few experimentally validated components of the glycolytic pathway in this organism (33). Their findings emphasized the potential importance of the enzyme in *Leptospira* metabolism and supported the broader notion that glucose-related pathways may play underappreciated roles in the bacterium’s physiology. Additional evidence for glucose responsiveness comes from studies of chemotaxis (34, 35). Chemotactic responses were evaluated for sugars, amino acids, fatty acids, and vitamins, and positive responses were observed for a subset of them, including glucose. Although chemotaxis is not directly linked to glucose transport or catabolism, the existence of a system capable of sensing or responding to glucose gradients is consistent with the possibility that glucose detection and acquisition have physiological relevance in *Leptospira*. Together with these earlier observations, our findings suggest that glucose-associated traits in *Leptospira* may be conditional rather than absent, becoming more apparent under specific physiological parameters such as host-like temperature.

Relatively small attention has been given to how temperature affects carbon source utilization in *Leptospira*. In contrast, numerous studies have examined temperature-dependent changes in *Leptospira* transcription and translation, motivated in part by the discrepancy between its optimal *in vitro* growth temperature and mammalian host body temperatures. Temperature is a critical regulator of gene expression and virulence in *Leptospira* species. Shifts from environmental temperatures (20–30 °C) to host-associated conditions (37–39 °C) induce substantial transcriptional reprogramming, affecting functional categories such as cell wall and membrane biogenesis, chemotaxis, and motility (2, 3). The expression of virulence factors, including multi-adhesive Lig proteins, is regulated by cis-acting RNA elements responsive to temperature changes (6), and further influenced by oxidative stress and host-specific factors (8). Strain-specific transcriptional profiles, such as those observed in *L. borgpetersenii* serovar Hardjo, suggest that temperature-mediated regulation may contribute to differences in clinical presentation and host colonization (25). Notably, while pathogenic *Leptospira* proliferates most actively at 30 °C, a temperature lower than mammalian host body temperature, they also encounter and persist under higher host-like temperatures (1). This raises the possibility that *Leptospira* exhibits distinct metabolic activities under environmental and host-like conditions. In the current study, we identified model-predicted metabolic changes under host-like temperature and *in vivo*–proximal condition, and experimentally supported their association with glucose uptake and glucose-supported growth *in vitro* (Figs 2, 3 and Fig. S3). Further investigation of temperature-dependent metabolic phenotypes during mammalian infection may inform future biomarker discovery and metabolic target-prioritization study for leptospirosis.

Public transcriptomic datasets involving temperature shifts (25, 27) and DMC conditions (28) were selected to contextualize the *Leptospira* metabolic model in this study. Of these, the L495 temperature model (Fig. S2) exhibited less pronounced differences compared to the other two models, JB197 temperature model and Fiocruz L1-130 DMC model (Fig. 2 and Fig. S3). This may be due in part to the different incubation conditions: the L495 temperature model was obtained relatively early at 18 hours after the start of the temperature switch, while the other data showed expression changes over a period of several days. The metabolic models used in this study still have room for improvement. In particular, the underlying transcriptomic data may not fully reflect metabolic steady-state conditions, an important assumption in constraint-based metabolic modeling, including flux balance analysis. Deviation from this assumption can limit the accuracy of predicted metabolic fluxes. Although the use of public data inevitably involves limitations such as experimental designs not optimized for the present question, incomplete or inconsistent metadata, and variability in data quality, these datasets can still provide useful insights when interpreted cautiously and integrated across multiple conditions. In this study, despite these limitations, the combined analyses served as a basis for hypothesis generation and increased confidence in the recurrent pattern of temperature-and host-like condition-associated glucose-related predictions.

*Leptospira* synthesize erythrose 4-phosphate from fructose 6-phosphate, which serves as a precursor for the biosynthesis of aromatic amino acids (12). These amino acids contribute to the structural stability of membrane proteins (36–38). They are also implicated in the functionality of several *Leptospira* virulence factors (39) and another bacterium such as *Listeria* (40). These considerations suggest that the predicted fructose 6-phosphate (F6P)-linked metabolic changes may have broader physiological consequences beyond growth, but their relevance to virulence-associated phenotypes remains to be tested.

This study has several limitations. First, because the aim of this study was to examine metabolic responses at host-like temperatures, the analyses and experiments focused primarily on pathogenic *Leptospira* strains associated with mammalian hosts. Therefore, it remains unclear whether the temperature-dependent glucose uptake and glucose-associated metabolic capacity identified here are traits specific to pathogenic *Leptospira* or represent a potential metabolic capacity more broadly conserved within the genus *Leptospira*. Future comparative analyses including nonpathogenic strains will be necessary to clarify how temperature-dependent carbon source utilization has evolved across the genus, while considering the ecological niches and optimal growth temperatures of each species. Second, this study did not directly trace the intracellular fate of glucose-derived carbon. Although experiments using fluorescent glucose analogues support temperature-dependent glucose uptake or accumulation, they do not define which metabolic pathways are utilized by imported glucose. Similarly, 2DG sensitivity is consistent with a contribution of glucose-associated processes to proliferation at 37°C, but it does not establish the essentiality of specific enzymatic reactions or pathways. Furthermore, although the upregulation of LA2133 and LB245 is consistent with the temperature-dependent uptake phenotype, to determine these gene products function as glucose transporters, it requires genetic or biochemical validation. Finally, because much of this study is based on *in vitro* experiments and transcriptome-integrated metabolic modeling, whether this phenotype operates during mammalian infection remains unresolved.

In conclusion, this study supports a model in which pathogenic *Leptospira* displays condition-dependent glucose uptake and glucose-associated metabolic activity that become apparent at host-like temperature, that is contrary to glucose-incompetent that has long been suggested. By combining transcriptome-integrated metabolic modeling with *in vitro* validation, we identified a temperature-dependent metabolic phenotype that refines the conventional view of *Leptospira* carbon metabolism. Future studies using stable glucose isotope tracing, comparative analyses across the genus, animal models, and clinical specimens will be required to define the relevance in biochemical, evolutionary, and *in vivo* aspects of the glucose-associated metabolic capacity.

## Methods

### Bacterial strain, culture conditions and reagents

*Leptospira interrogans* serovar Manilae strain L495 was used in this study. L495 was cultured in EMJH medium (BD Biosciences, Franklin Lakes, NJ, USA) at either 30 °C or 37 °C under static conditions. Bacterial counts were determined using a Petroff-Hausser counting chamber (Sunlead Glass, Saitama, Japan) under a dark-field microscope (BX-53, Olympus, Tokyo, Japan). Other reagents used in this study were as follows: 2-Deoxy-D-glucose (Cat# 10722-11, Nacalai Tesque Inc., Kyoto, Japan)

### Genome-scale metabolic network reconstruction in pathogenic *Leptospira*

Genome-scale metabolic models of pathogenic *Leptospira* were constructed using Reconstructor version 1.0.76 (23). Amino acid sequence data annotated with PATRIC (41) used as input were obtained from the Bacterial and Viral Bioinformatics Resource Center (BV-BRC 3.47.8) (42). During model reconstruction, both EMJH medium and the complete medium required by Reconstructor were specified (Table S2). All reaction identifiers in the resulting models were derived from the ModelSEED database (43).

### Integration of public transcriptome data to contextualize GENREs

Reconstructed GENREs were integrated and contextualized using publicly available transcriptomic data with the RIPTiDe algorithm (26). Flux balance analysis (FBA) was performed within RIPTiDe by running 1,000 simulations with random sampling, and the resulting flux distributions were compared and analyzed. The following RNA-seq datasets were used: (i) GSE168507 (25), *L. borgpetersenii* serovar Hardjo strain JB197 incubated at 29 °C and 37 °C; (ii) GSE92976 (27), *L. interrogans* serovar Manilae strain L495 cultured at 30 °C and 37 °C; and (iii) GSE53818 (28), *L. interrogans* serovar Copenhageni cultured in a dialysis membrane chamber (DMC) model embedded in the rat peritoneal cavity or *in vitro* at 30 °C. All RNA-seq data used in this study were obtained from previously published datasets and were not reprocessed from raw sequencing files. Important reactions were selected based on random forest feature importance, expressed as the mean decrease in accuracy, as described in a previous study (44). A larger mean decrease in accuracy indicates that permutation of the corresponding reaction has a greater negative effect on classification performance.

### Fluorescent-labeled glucose uptake assay

Glucose uptake by *Leptospira* was quantified using the Glucose Uptake Assay Kit-Green (UP02, Dojindo, Kumamoto, Japan), following the manufacturer’s instructions. L495 strain was incubated for 48 hours in medium either standard EMJH-standard or - supplemented with 10% glucose prior to the assay. Fluorescence-conjugated glucose uptake was measured using a SPARK 10M microplate reader (TECAN, Männedorf, Switzerland) in fluorescence detection mode (excitation, 470 nm; emission, 520 nm). Fluorescence intensity was normalized to bacterial cell number and expressed as relative uptake.

### Growth curve assay and fitting

Bacterial growth was monitored using an iMark microplate reader (Bio-Rad, Hercules, CA, USA) by measuring absorbance at 450 nm. For glucose supplementation experiments, EMJH medium was supplemented with glucose. For growth inhibition assays, 2-deoxy-D-glucose (2DG) was added to EMJH. Growth curves were generated based on absorbance data using the growthcurver package in R (version 0.3.1, (45)). Based on a logistic growth model, the area under curve (AUC) was estimated.

### Gene expression analysis

*L. interrogans* strain L495 was cultured in EMJH medium at 30 °C and 37 °C for 24 and 48 hours. Cells were harvested by centrifugation, and total RNA was extracted using the Direct-zol™ RNA Microprep kit (R2063, Zymo Research Corp., Irvine, CA, USA). cDNA was synthesized using the ReverTra Ace qPCR RT Master Mix with gDNA Remover (FSQ-301, TOYOBO Co., Ltd., Osaka, Japan). Quantitative real-time PCR was performed using the CFX Connect Real-Time PCR Detection System (Bio-Rad, Hercules, CA, USA) with Thermo Scientific SYBR Green qPCR Master Mix (A66732, Thermo Fischer Scientific Inc., Waltham, MA, USA). Gene expression levels were quantified using the ΔΔCt method, with 16S rRNA as the internal reference gene. Primers used in this study are listed in Table S3.

## Statistical analysis

All statistical analyses in this study including the Welch’s t-test, one-way ANOVA followed by Tukey’s HSD test, Wilcoxon rank-sum test, Dunn’s test with Bonferroni correction, and permutational ANOVA (PERMANOVA) were performed using R (version 4.4.0). Normality was assessed using the Shapiro–Wilk test or Q–Q plots prior to applying parametric tests. Statistical significance was defined as *p* < 0.05.

## Data availability

All code used in this study, the generated genome-scale metabolic models, and the raw and processed data supporting the findings are available in a GitHub repository (https://github.com/O2U/Lepto-Gluc-Metab).

## Acknowledgement

We thank the members of the Department of Microbiology and Immunology, Faculty of Medicine, Fukuoka University for helpful discussions and technical support. This work was supported by JSPS KAKENHI (Grant No. 24K10225 to R.O. and 24K11646 to M.Y.), funding from Fukuoka University (Grant No. 247302 and GR2613) to R.O., AMED/NIH/CRDF (Grant No. JP23jk0210044 to F.O. and G.L.K.), AMED/NIH/ISTC (Grant No. JP25jk0210054 to R.O. and J.A.P.), Takeda Science Foundation to R.O., and Heisei Memorial Research Fund to R.O. The funders had no role in study design, data collection and analysis, decision to publish, or preparation of the manuscript.

## Author contribution

R.O., M.Y., F.O., J.A.P. G.L.K. and K.H. conceived the study and designed the experiments. R.O., D.A.P. performed the genome-scale metabolic network reconstruction and flux analysis. R.O., M.Y., F.O. and T.S. performed the experiments and collected data. R.O. performed statistical analyses and drafted the paper and figures. All authors reviewed and approved the final manuscript

